# Recently formed context fear memories can be retrieved without the hippocampus

**DOI:** 10.1101/843342

**Authors:** Jamie N. Krueger, Jacob H. Wilmot, Yusuke Teratani-Ota, Kyle R. Puhger, Sonya E. Nemes, Marrisa Lafreniere, Brian J. Wiltgen

## Abstract

The current study determined if inactivation of the dorsal hippocampus impairs the retrieval of newly formed context fear memories. This region was silencing by activating inhibitory neurons or by hyperpolarizing pyramidal cells directly. When inhibitory neurons were stimulated with ChR2, memory retrieval was significantly impaired. In contrast, when the same neurons were activated with the excitatory DREADD hM3Dq, retrieval was not affected. This dissociation was not due to differences in inhibition, as both manipulations activated interneurons and reduced excitation throughout the dorsal hippocampus. Therefore, we hypothesize that the retrieval deficit caused by ChR2 stimulation is due to an immediate reduction in hippocampal activity that does not provide enough time for other brain regions to compensate. Stimulation of DREADDs, on the other hand, produces a gradual loss of excitation that takes several minutes to reach asymptote. This appears to be a sufficient amount of time for extra-hippocampal structures to become engaged and express context fear. Implications for theories of hippocampal function, systems consolidation and memory retrieval are discussed.

## Introduction

Encoding and retrieving new context fear memories depends on the hippocampus [1–4]. In contrast, older memories can often be retrieved without this structure [5–7]. This difference is thought to result from systems consolidation, a process that gradually transfers memories from the hippocampus to the cortex [8,9]. In order for systems consolidation to occur, a series of coordinated events have to take place after learning: sharp-wave ripple oscillations in the hippocampus need to be synchronized with sleep spindles in the cortex [10]; immediate early gene expression has to be induced and maintained in cortical neurons during sleep [11–14]; and cortical synapses have to be potentiated so they can respond to retrieval cues during remote memory testing. Consequently, the existence of systems consolidation and the speed at which it occurs, varies markedly across studies [6,7,12,15].

Retrieving remote memories with the cortex also requires a number of events to occur. The hippocampus has to become less responsive to retrieval cues, so it is not active during testing [8,12,16]. Neurons in the prefrontal cortex (PFC) and the anterior cingulate cortex (ACC) have to grow new spines after learning so context fear can be expressed several weeks later [5,9,17]. And, if the hippocampus has been silenced experimentally, cortical regions needs several minutes to recover before they can successfully retrieve remote memory. When animals are tested immediately after hippocampal inactivation, context fear cannot be expressed [18]. These data illustrate that remote memory retrieval results from dynamic interactions between the hippocampus and cortex that take place continually after learning [19,20].

New context memories can also be retrieved without the hippocampus in some situations. For example, if the hippocampus is damaged prior to training, the PFC is able to acquire and express context fear [4,21–23]. Recent memories can also be retrieved if the hippocampus is silenced pharmacologically for a prolonged period prior to testing [24–26]. These data indicate that, under some conditions, the cortex appears to play a similar role in the expression of recent and remote context fear.

In the current study, we compared the effects of acute and prolonged silencing of the hippocampus on the expression of recently acquired context fear. Similar to remote memory, we hypothesized that acute optogenetic silencing would produce larger deficits than prolonged inactivation with DREADDs. We also examined the impact of inhibition produced by interneuron stimulation to that observed when pyramidal cells were hyperpolarized directly. Consistent with our prediction, acute silencing of the hippocampus led to memory impairments while prolonged inactivation did not. Stimulation of inhibitory neurons also produced more extensive inhibition than direct hyperpolarization of pyramidal cells, as recently reported [27]. Unexpectedly, some inhibitory manipulations (e.g. CaMKII-ArchT and hSyn-hM4Di stimulation) led to widespread increases in hippocampal activity that profoundly impaired memory retrieval. We suspect these changes were produced by local disinhibition that spread throughout the network [28,29].

These results demonstrate that the dorsal hippocampus is not required for the retrieval of recently acquired context fear. This finding is inconsistent with models of systems consolidation that assume a lengthy storage process is required for memories to become independent of this region. However, if the hippocampus goes offline rapidly or becomes hyperactive, then recently formed context fear memories cannot be expressed. Implications for theories of hippocampal function, memory consolidation and retrieval are discussed.

## Results

### Acute, but not prolonged activation of inhibitory neurons impairs memory retrieval

It was recently shown that activation of GABAergic neurons produces more efficient and widespread inhibition of pyramidal cells than direct photoinhibition using light-gated ion pumps [27,30]. Given that the dorsal hippocampus is a large structure, we first silenced it by stimulating inhibitory neurons with ChR2 or hM3Dq. To do this, we used the Dlx enhancer, which produces expression in forebrain GABA-ergic neurons (**Figure 1A**) [31]. Using the mDlx enhancer, we targeted the excitatory opsin ChR2 to inhibitory neurons within the CA1 region of the dorsal hippocampus (**Figure 1B**). We also used the hDlx enhancer to target the excitatory DREADD hM3Dq to the same brain region (**Figure 1C**). Viral expression was limited to inhibitory interneurons as previously described [31].

**Figure 1.**
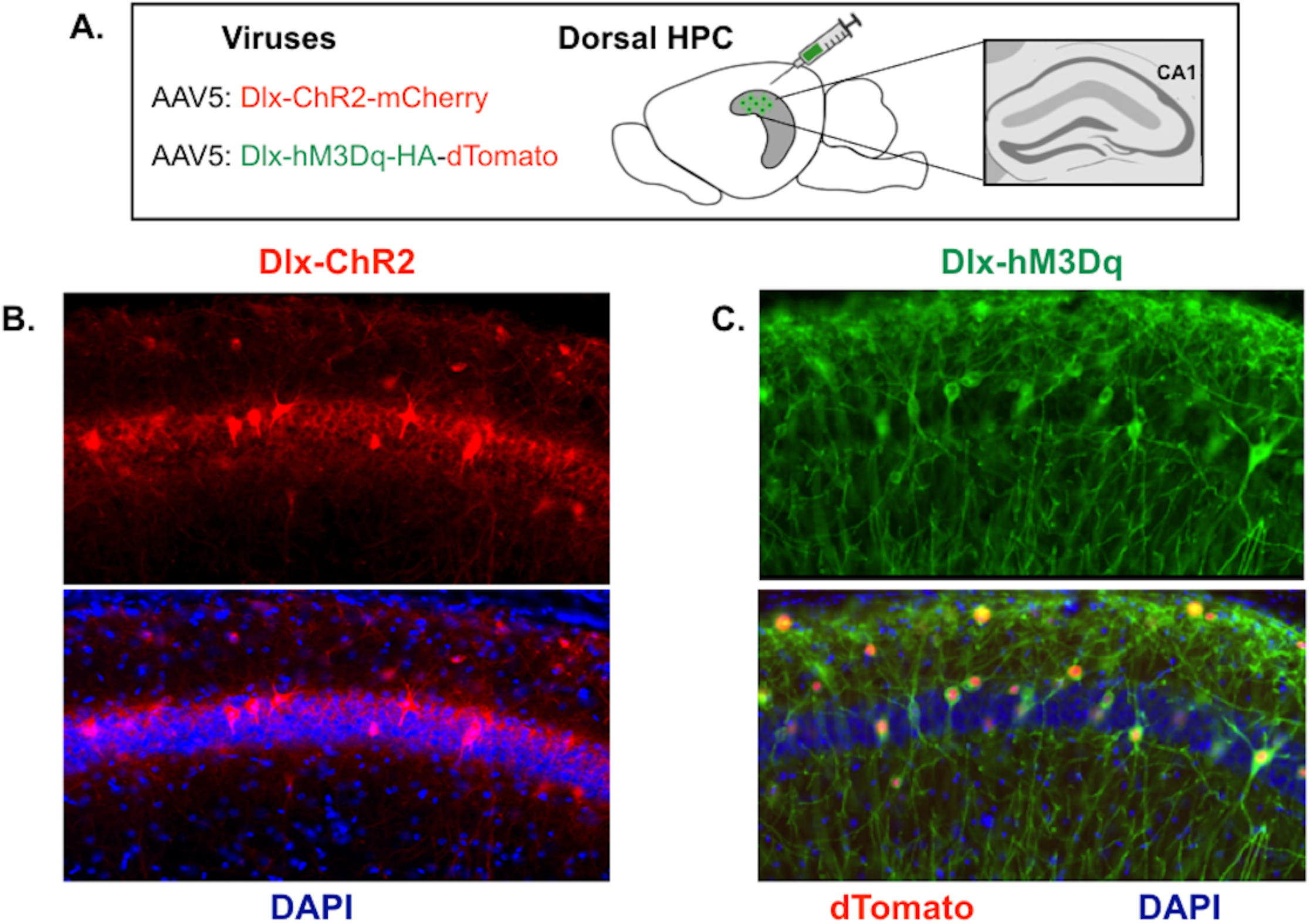
Expression of ChR2 and hM3Dq in inhibitory neurons. A. Dlx-ChR2-mCherry or Dlx-hM3Dq-dTomato were expressed in CA1 of the dorsal hippocampus (image adapted from [57]. B. *Left top*: Dlx-ChR2 expression (red) in dorsal CA1. *Left bottom*: Dlx-ChR2 expression merged with DAPI (blue) in dorsal CA1. C. *Right top*: Dlx-hM3Dq HA-tag expression (green) in dorsal CA1. *Right bottom*: Dlx-hM3Dq HA-tag expression (green) and Dlx-hM3Dq dTomato nuclear expression (red) merged with DAPI (blue) in dorsal CA1.

We first examined the effects of acute activation of inhibitory neurons on memory retrieval. Two weeks following Dlx-ChR2 infusion (**Figure 2A)**, mice were trained on contextual fear conditioning. One day later, the animals were returned to the chamber for a memory test (**Figure 2B**). Following a 3-minute laser off period, animals underwent 3 minutes of laser stimulation (473nm, 20Hz, 10mW). This was repeated once. Stimulation of inhibitory neurons impaired memory retrieval in Dlx-ChR2 mice (n=10) compared to controls (n=6) during the laser on period (**Figure 2C**) (Group x laser interaction F (1,15), p =.0012, Fisher’s LSD post-hoc tests, Laser off (p =.54), Laser on (p = .0064)). We next determined if ChR2 stimulation activated interneurons and inhibited pyramidal cell activity using c-Fos expression. Given that our optogenetic testing parameters included both laser off and laser on periods (which could obscure reductions in activity as measured by c-Fos expression), we conducted a second test the following day where ChR2 was stimulated for 12 minutes. As expected, we found that pyramidal cell activity was significantly reduced in Dlx-ChR2 animals (**Figure 2D**) (t=7.366, df = 13, p < 0.0001). We also found that interneuron activity was significantly increased in Dlx-ChR2 animals (**Figure 2E**) (t=4.226, df=13, p = .0010). These data indicate that activation of inhibitory neurons in dCA1 reduces the activity of pyramidal neurons and impairs memory retrieval.

**Figure 2.**
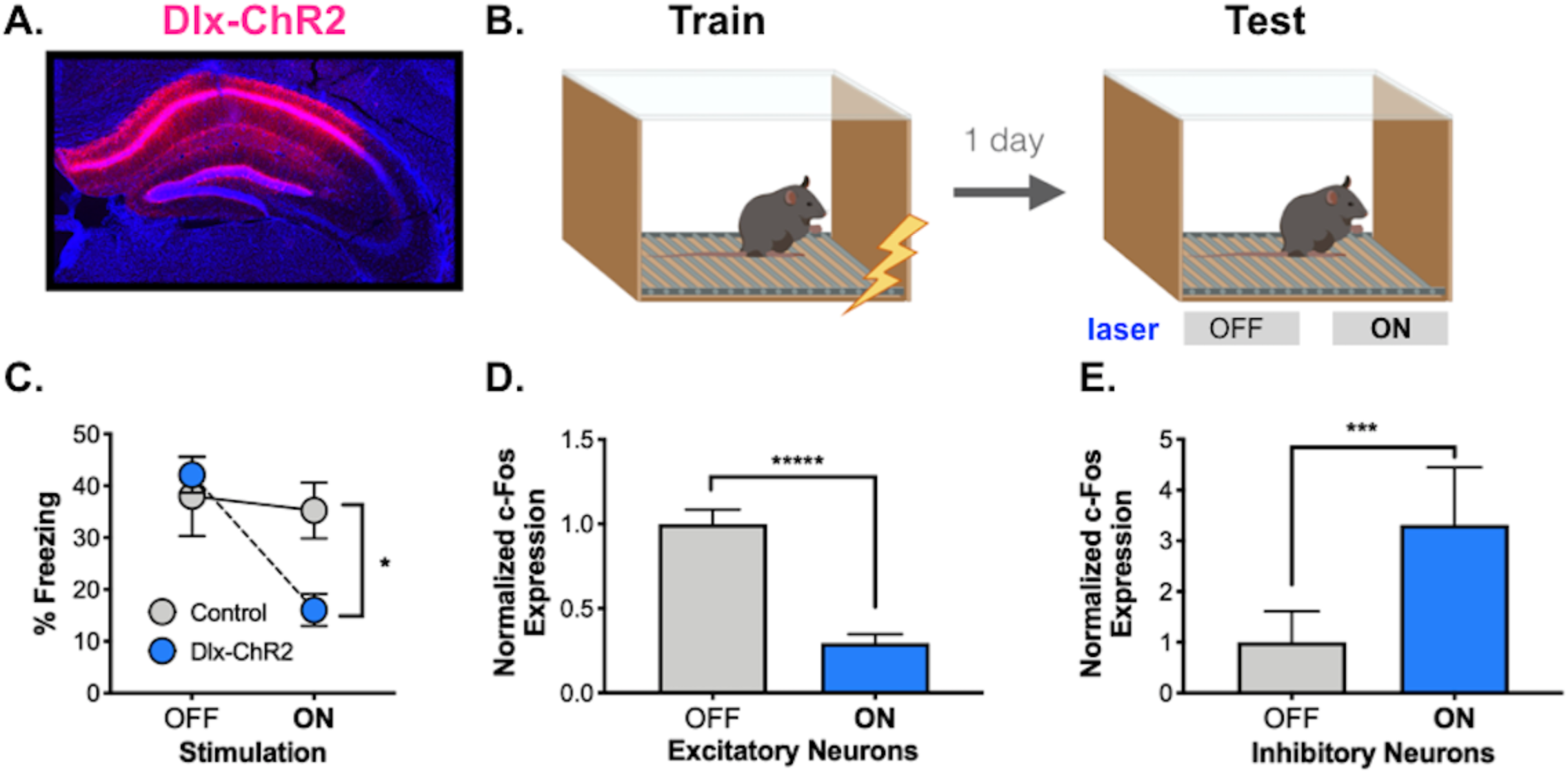
Acute activation of inhibitory neurons. A. Dlx-ChR2 expression (pink) in the dorsal hippocampus. B. Behavioral paradigm. Dlx-ChR2 animals were trained in contextual fear conditioning (4 × .75mA shocks). One day later animals were returned to the testing chamber for a 12-minute test. Animals received a 3-minute baseline, then 3 minutes of blue laser (473nm, 20Hz, 10mW) stimulation. This was repeated once. C. Dlx-ChR2 animals (blue) froze less during Laser ON periods than controls (gray). D. c-Fos expression was lower in excitatory pyramidal neurons in dCA1 of Dlx-ChR2 (blue) vs. control (gray) animals. E. c-Fos expression was higher in inhibitory neurons in dCA1 of Dlx-ChR2 (blue) vs. control (gray) animals. All data are expressed as mean +/- SEM.

Next, we examined the effects of prolonged activation of inhibitory neurons on memory retrieval using the excitatory DREADD hM3Dq (**Figure 3A**). As in the previous experiment, Dlx-hM3Dq mice were trained on contextual fear conditioning and then tested 1-day later. One hour prior to re-exposure to the context, animals were given an injection of 5mg/kg CNO (n=10) or vehicle (n=9) (**Figure 3B**). In contrast to acute activation of inhibitory neurons, prolonged stimulation with CNO did not impair memory relative to vehicle-treated animals (**Figure 3C**) (No effect of group F (1,17) = 1.931, p =.1826). Examination of c-Fos expression after testing, revealed that similar to our Dlx-ChR2 data, stimulation of Dlx-hM3Dq expressing neurons decreased the activity of pyramidal cells (**Figure 3D**) (t=11.68, df=17, p<0.0001) and increased the activity of inhibitory neurons (**Figure 3E**) (t=11.44, df=17, p<0.0001). Therefore, when the dorsal hippocampus is inactivated for a prolonged period, newly formed context fear memories can be retrieved by other brain regions. These results are similar to the effects of acute and prolonged hippocampal silencing during remote memory retrieval [18].

**Figure 3.**
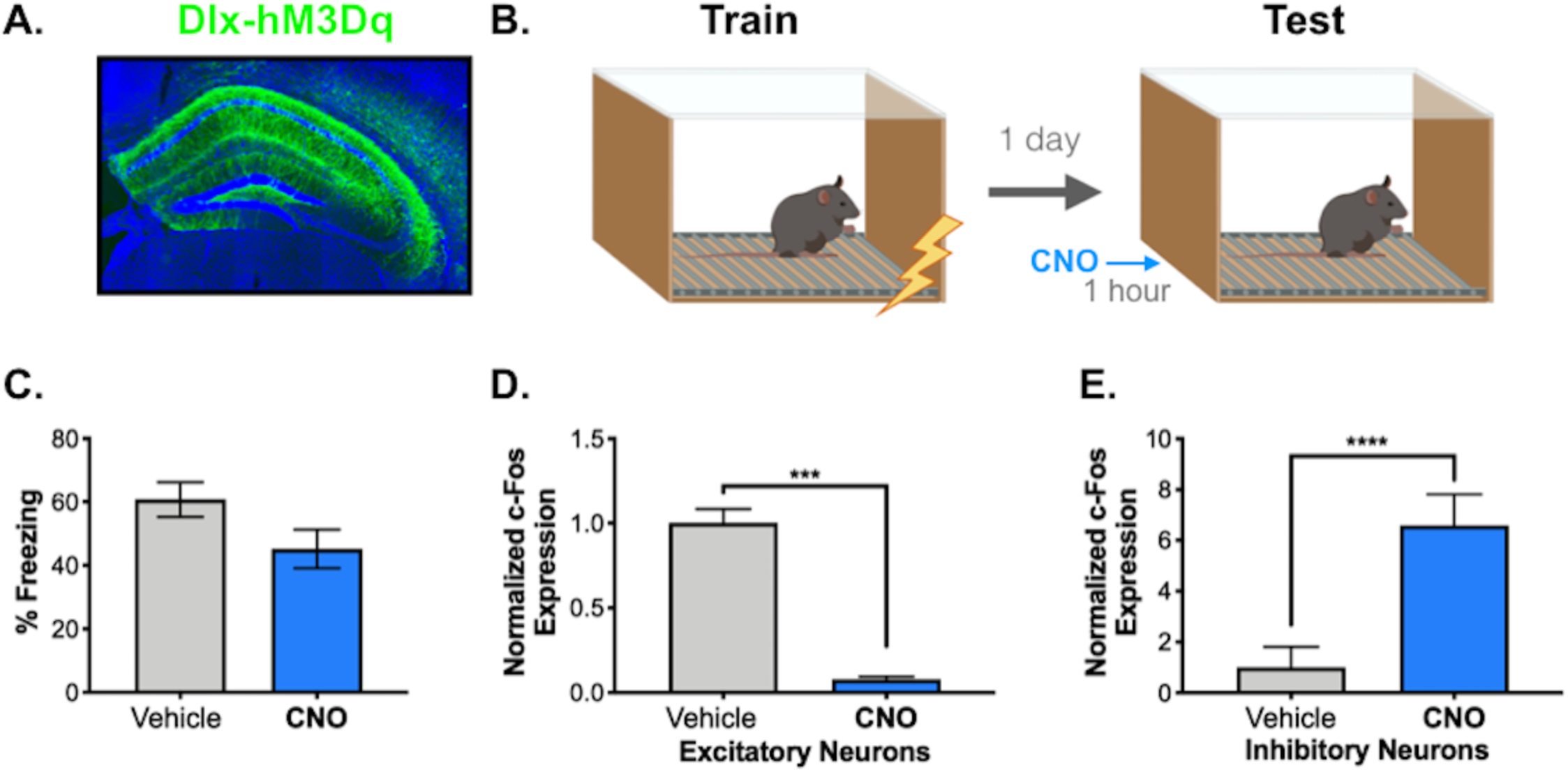
Prolonged activation of inhibitory neurons. A. Dlx-hM3Dq expression (green, HA tag) in the dorsal hippocampus. B. Behavioral paradigm. Dlx-hM3Dq animals were trained in contextual fear conditioning (3 × .4mA shocks). One day later, animals were returned to the chamber for a 12-minute memory test following a 5mg/kg CNO or vehicle IP injection. C. There was no difference in freezing over the 12-minute testing period between the vehicle (gray) and CNO-treated (blue) animals. D. c-Fos expression was lower in excitatory pyramidal neurons in dCA1 of CNO-treated (blue) vs. vehicle-treated (gray) animals. E. c-Fos expression was increased in inhibitory neurons in dCA1 of CNO-treated (blue) vs. vehicle-treated (gray) animals. All data are expressed as mean +/- SEM.

Stimulation of GABAergic neurons with ChR2 can produce inhibition that extends 1-2mm from the center of the optic fiber [27,30]. To determine the extent of GABAergic stimulation in our experiments, we quantified c-Fos expression in inhibitory neurons at multiple anteroposterior (AP) coordinates in the dorsal, intermediate and ventral hippocampus in a subset of animals from each group (ChR2 n=5, control n=5; hM3Dq CNO n=4, vehicle n=4). To avoid the accidental inclusion of excitatory neurons, quantification was restricted to the SO, SR and SLM cell layers, where the majority of inhibitory neurons are found (**Figure 4A**). We observed that ChR2 stimulation increased c-Fos expression in inhibitory neurons compared to control animals throughout the entire extent of the dorsal hippocampus (**Figure 4B**) (Group x AP coordinate interaction, F (2,16) = 6.12, p = .0106). Similar to published work, this increase extended at least .8 mm AP from the fiber tip (Fisher’s LSD post-hoc tests, Anterior (p = .042), Intermediate (p = .0048)). Elevated c-Fos expression in inhibitory neurons was not observed in the ventral hippocampus (Fisher’s LSD post-hoc test, Posterior (p = .3294)) [27,30]. A similar spatial profile was found when GABAergic neurons were activated with hM3Dq (**Figure 4C**) (Group x AP coordinate interaction, F (2,12) = 42.2, p = <.0001; Fisher’s LSD post-hoc tests, Anterior (p <.0001), Intermediate (p=.0003), Posterior (p=.1060)). These data verify that both manipulations activated inhibitory neurons throughout the dHPC.

**Figure 4.**
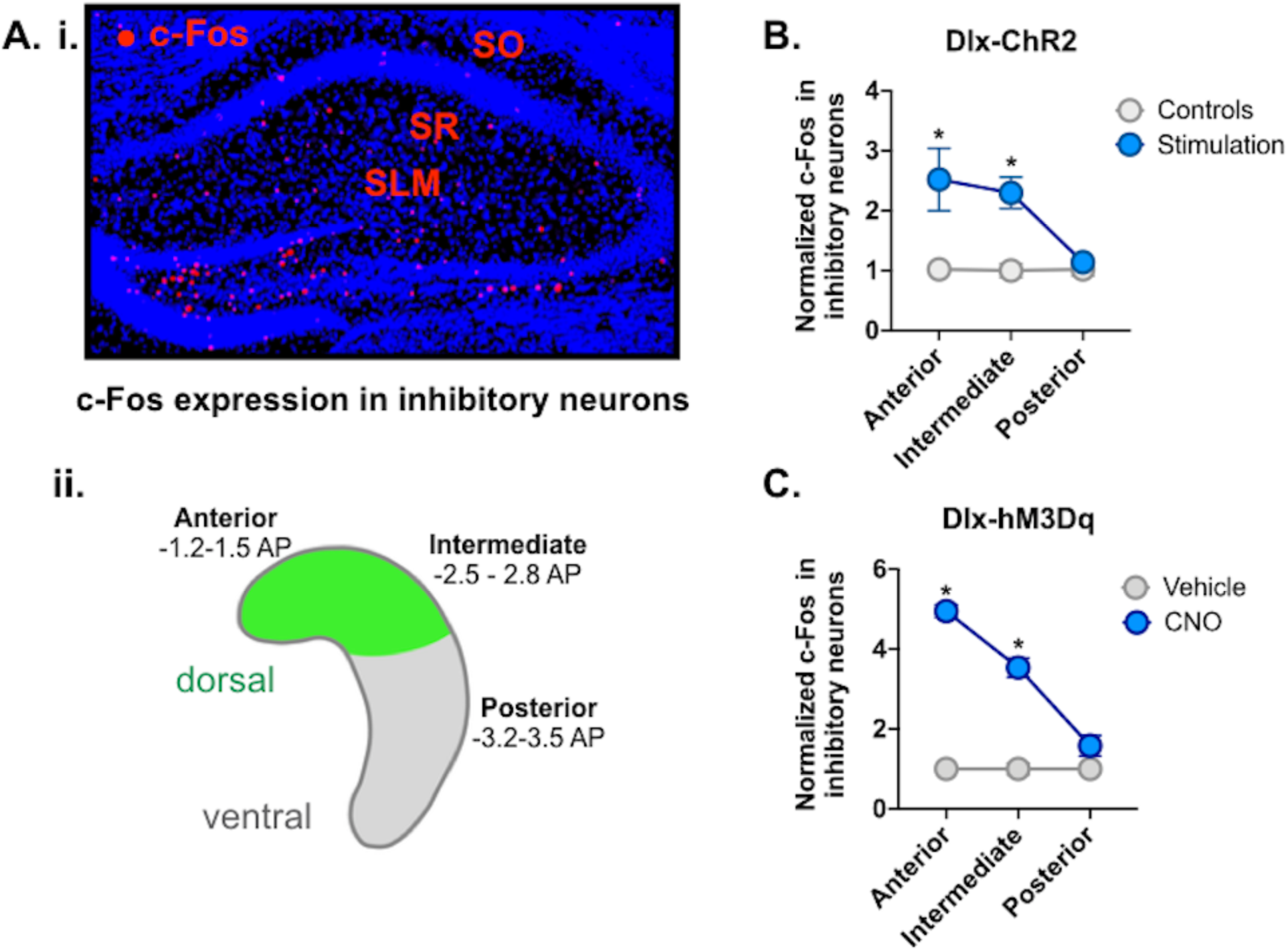
Widespread activation of inhibitory neurons in the dorsal hippocampus. A. i. Anterior dHPC of representative CNO-treated Dlx-hM3Dq animal showing c-Fos expression (red) in inhibitory strata: stratum oriens (SO), stratum radiatum (SR), stratum lacunosum-moleculare (SLM). A. ii. Schematic of HPC showing location of virus expression (green) in dorsal HPC. Representative slices were taken for c-Fos analysis from anterior (−1.2 to −1.5 AP), intermediate (−2.5 to −2.8 AP) and posterior/ventral HPC (−3.2 to −3.5 AP). B. c-Fos expression in inhibitory neurons of anterior and intermediate HPC was increased in Dlx-ChR2 laser-stimulated animals (blue). There was no difference in c-Fos expression in posterior HPC. C. c-Fos expression in inhibitory neurons of anterior and intermediate hippocampus was increased in Dlx-hM3Dq CNO-treated animals (blue). There was no difference in c-Fos expression in posterior HPC. All data are expressed as mean +/- SEM.

### Acute and prolonged activation of pyramidal neurons impair memory retrieval

Next, we compared the effects of acute and prolonged activation of pyramidal neurons during memory retrieval (**Figure 5A**). In the first experiment, the excitatory DREADD hM3Dq was expressed in dCA1 excitatory neurons using the CaMKII promoter. Similar to our previous DREADD experiment, mice were trained in contextual fear conditioning and memory was tested 24 hours later one hour after an IP injection of .5mg/kg CNO (n=5) or vehicle (n=5). We found that prolonged stimulation of excitatory neurons profoundly impaired memory retrieval (**Figure 5B**) (Main effect of group, F (1, 8) = 17.9, p = 0.003). During the test, there was also a significant group x time interaction (F (3, 24) = 9.39, p = .0003) indicating that controls animals extinguished while the CNO group did not. To confirm that activation of hM3Dq increased the activity of pyramidal neurons during the context test, we quantified c-Fos expression in dCA1 (**Figure 5C**). As expected, there was a significant increase in c-Fos labeling in CNO mice compared to the vehicle-treated group (t (8) = 12.13, p < 0.0001).

**Figure 5.**
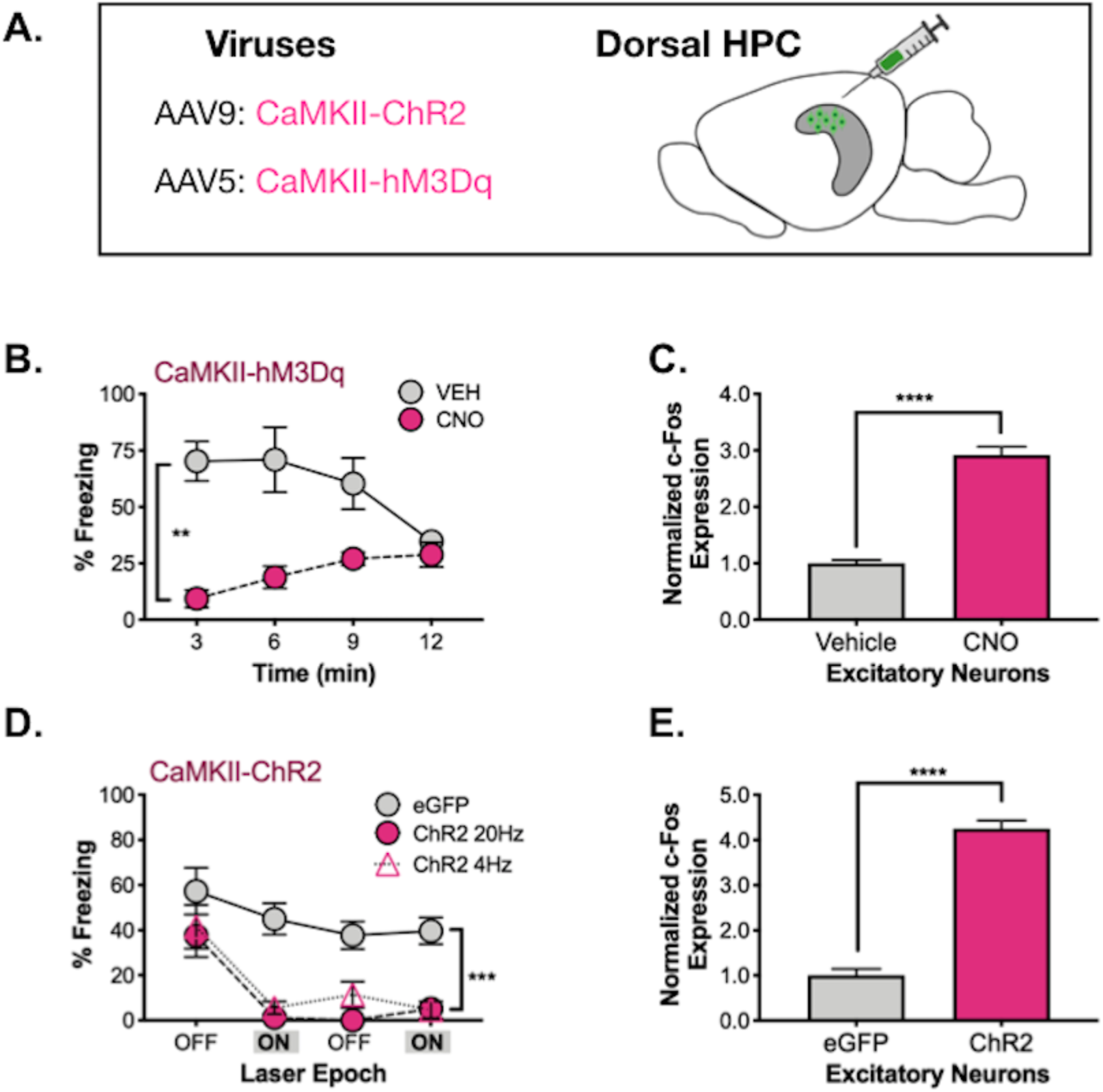
Prolonged vs. acute activation of excitatory neurons. A. CaMKII-ChR2 or CaMKII-hM3Dq were infused into dHPC. B. During memory testing, hM3Dq CNO-treated animals (pink) froze less than their vehicle-treated counterparts (gray). C. c-Fos expression in excitatory neurons of dCA1 was elevated in hM3Dq CNO-treated animals (pink) compared to their vehicle-treated counterparts (gray). D. During memory testing, ChR2 (20Hz – pink circles, 4Hz – pink triangles) animals and eGFP animals (gray) did not differ during the first 3-minute laser OFF epoch. Following laser stimulation, 20 and 4Hz stimulated animals froze less than controls over the remainder of the testing period. E. c-Fos expression was elevated in excitatory neurons of dCA1 in ChR2 20Hz-stimulated animals (pink) compared to controls (gray). All data are represented as mean +/- SEM.

In order to acutely activate pyramidal neurons, we expressed ChR2 in excitatory neurons using the CaMKII promoter (**Figure 5A**). As in the previous experiment, mice were trained in contextual fear conditioning, then 24 hours later were returned to the chamber for a retrieval test. Following a 3-minute laser off period, animals received 3 minutes of laser stimulation (473nm, 20Hz, 10mW). This sequence was repeated one time. Excitatory neurons were stimulated at either 4Hz (n = 6) or 20Hz (n = 5) because the former induces freezing in CA1 “engram cells” while the latter does not [32]. However, we did not observe a difference between these stimulation frequencies, so the groups were combined for statistical analyses. Following the baseline period, laser stimulation significantly decreased freezing in ChR2 animals but not in control mice (**Figure 5C**) (Stimulation period x Group interaction F (1, 15) = 5.938, p = .0278), Fisher’s LSD post-hoc tests, ChR2 (p < .0001), eGFP (p = .1386). Interestingly, freezing remained low in ChR2 mice after the laser turned off and did not recover for the remainder of the test session (Main effect of group F (1, 15) = 36.45, p < .0001, No effect of stimulation period F (1, 15) = .024, p = .8786, No group x stimulation period interaction F (1, 15) = .608, p = .4476). As expected, analysis of c-Fos expression found increased activation of dCA1 neurons in ChR2 mice compared to controls (t (8) = 14.23, p<0.0001) (**Figure 5D**). Together, these results demonstrate that artificial increases in excitatory activity in the dHPC severely impairs the expression of context fear. In contrast to inhibition, deficits were observed following acute and prolonged increases in excitation.

In the next experiment, we inhibited excitatory neurons directly and examined the effects on memory retrieval. To do this, we expressed the inhibitory opsins ArchT (n = 6) and NpHR (n = 6) or the inhibitory DREADD hM4Di in excitatory neurons using the CaMKII promoter (**Figure 6A**). For the optogenetic manipulations, mice were trained and tested as described above, except that yellow (561nM, 15mW) or green (532nM, 10mW) light was delivered during the stimulation periods. DREADD expressing animals received an IP injection of 5mg/kg CNO (n=7) or vehicle (n=6) one hour prior to a 12-minute test.

**Figure 6.**
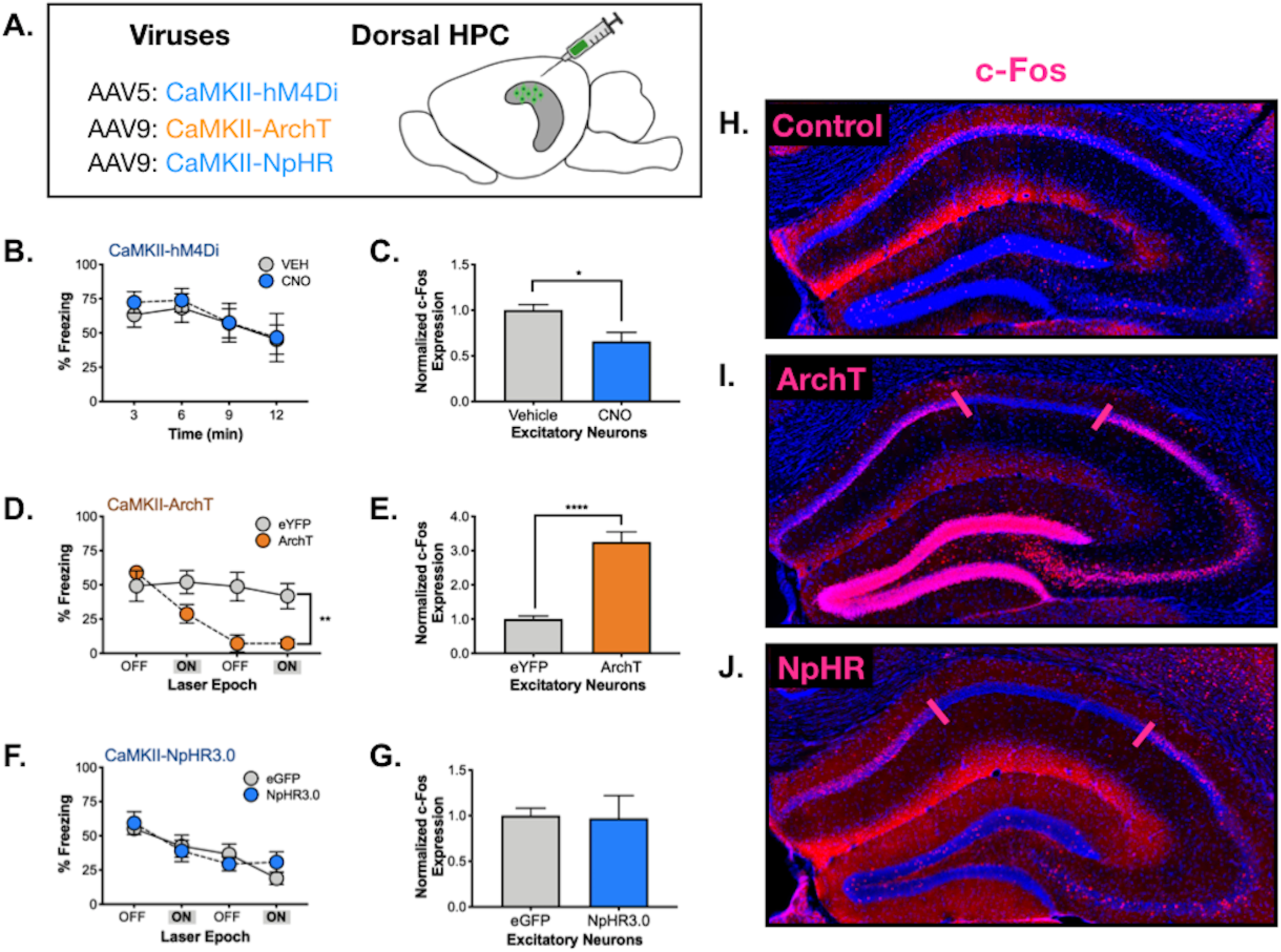
Prolonged vs. acute inhibition of excitatory neurons. A. CaMKII-hM4Di, CaMKII-ArchT or CaMKII-NpHR were infused into the dHPC in order to silence excitatory neurons. B. There was no difference in freezing during testing between hM4Di CNO (blue) and vehicle-treated (gray) animals. C. There was a small, but significant decrease in dCA1 c-Fos expression in CNO-treated hM4Di expressing animals (blue). D. There was no difference in freezing during the initial 3-minute laser OFF period between ArchT (orange) and eYFP (gray) expressing animals. Following the first laser ON epoch, ArchT animals froze less than controls over the remainder of the testing period. E. There was a significant **increase** in dCA1 c-Fos expression in ArchT animals (orange) compared to controls. F. There was no difference in freezing during laser OFF or ON epochs in NpHR vs. control animals. G. There was no difference in dCA1 c-Fos expression between NpHR and control animals. H. c-Fos expression (pink) in dHPC of a control animal. I. c-Fos expression (pink) in an ArchT-expressing animal. There is no c-Fos expression underneath the laser (denoted between pink lines), but expression is elevated throughout dentate gyrus, CA3 and CA1 surrounding the area of silencing. J. c-Fos expression (pink) in dHPC of a NpHR-expressing animal. c-Fos is absent in the area underneath the laser (denoted between pink lines), but it not elevated outside of this zone. Data are represented as mean +/- SEM.

Similar to our Dlx-hM3Dq data, prolonged silencing with CaMKII-hM4Di did not impair memory retrieval (No effect of treatment, F (1, 11) = 0.077, p = 0.785; Main effect of time F (3, 33) = 6.98, p = .0009, No treatment x time interaction F (3, 33) = .216, p = .8844) (**Figure 6B).** Analysis of c-Fos expression after the test revealed that hM4Di stimulation reduced the activity of CA1 pyramidal neurons although the size of this decrease was quite modest (t (10) =2.95, p = 0.01) (**Figure 6C**).

In contrast to prolonged silencing, acute inactivation of the dHPC with ArchT profoundly impaired memory retrieval (**Figure 6D**). Interestingly, the behavioral deficit looked similar to that observed in our CaMKII-ChR2 experiment; laser stimulation reduced freezing in ArchT animals after the baseline period (Group x Stimulation period interaction F (1, 9) = 9.254, p = .014, Fisher’s LSD post-hoc tests, Laser off (p .384) Laser on (p = .049) and it did not recover for the remainder of the session (Main effect of group F (1, 9) = 14.39, p = .0043, No effect of Stimulation period F (1,9) = 1.476, p = .2554), No Group x Stimulation period interaction F (1, 9) 1.539, p = .2461). Given this fact, we examined c-Fos expression 90-minutes after the testing session and found that ArchT stimulation produced in an unexpected increase in excitatory activity compared to controls (n = 5) rather than a decrease (t (8) = 5.92, p=0.0004) (**Figure 6E**).

To determine if this effect was selective to ArchT, we next stimulated NpHR during testing. Surprisingly, this manipulation had no effect on memory retrieval (No effect of group, F (1,9) = 0.05, p = 0.8182; Main effect of time, F (3, 27) = 9.23, p = 0.0002; No group x stimulus period interaction, F (3,27) = 0.8, p = 0.47) (**Figure 6F**) and did not alter c-Fos expression relative to controls (n = 6) (t (7) = 0.11, p = 0.92) (**Figure 6G**). However, as noted above, our optogenetic tests contain both laser off and on periods, which could obscure inhibitory effects on pyramidal cell activity. In addition, rebound excitation has been observed after optogenetic silencing, which could elevate c-Fos expression in the ArchT and NpHR groups. Therefore, we conducted a longer test session (25 min) during which these inhibitory opsins were stimulated continuously **(Figure 6)**. Compared to controls (**Figure 6H**), we observed that stimulation of ArchT and NpHR reduced c-Fos expression ≈1mm around the tip of the optic fiber (**Figure 6I,J**). This decrease was not observed in control animals, suggesting that it was not caused by laser stimulation alone. In addition to decreasing activity near the fiber tip, ArchT also produced a large increase in c-Fos expression outside of this region that extended into CA3 and the dentate gyrus. A similar increase was not observed following NpHR stimulation. Given that the silenced area was similar for both opsins, we hypothesize that CaMKII-ArchT stimulation impaired memory retrieval because it increased excitatory activity in the dHPC.

ArchT is a proton pump and stimulating it from extended periods (< 1 min) can raise intracellular pH in synaptic terminals and increase glutamate release [33]. Stimulation can also decrease extracellular pH and induce action potentials in neighboring neurons by activating acid-sensing ion channels (ASICs) [34,35]. Given that these excitatory effects are not observed with brief stimulation, we examined freezing during the context test in 20s intervals (**Figure S1A**). We also compared the CaMKII-ArchT data to the behavioral deficit observed when inhibitory neurons were stimulated with ChR2. Despite the fact that both of these manipulations produce rapid inactivation of pyramidal cells [27], only Dlx-ChR2 stimulation produced immediate decreases in freezing **(Figure S1BB)**. Decreases did not emerge in the CaMKII-ArchT group until CA1 had been stimulated for ≈100-140s (**Figure S1A**). When laser stimulation terminated, freezing returned to control levels in the Dlx-ChR2 group but did not recover in ArchT animals. These data strongly suggest that stimulation of CaMKII-ArchT impairs memory retrieval because it increases excitatory activity in the dHPC.

### Prolonged inhibition of both excitatory and inhibitory neurons

In a recent study, hM4Di was expressed in the dHPC under control of the neuron-specific promoter hSyn and used to hyperpolarize cells during object recognition tests [29]. The authors found that stimulation impaired memory retrieval and produced an unexpected increase in excitatory activity. Follow-up experiments revealed that the increase in activity resulted from reduced GABA’ergic tone caused by hyperpolarization of inhibitory neurons. To determine if similar effects would be observed with context fear conditioning, we expressed hM4Di in dorsal hippocampal neurons using the hSyn promoter. Similar to the previous report, CNO administration prior to testing significantly impaired the retrieval of context fear and increased c-Fos expression in pyramidal neurons (**Figure S1C-E**). Therefore, similar to our CaMKII-ArchT data, these results highlight the importance of understanding how network activity is affected by targeted optogenetic and chemogenetic manipulations.

## Discussion

In the current study, we determined if inactivation of the dorsal hippocampus impairs the retrieval of newly formed context fear memories. We silenced this structure by activating inhibitory neurons or by hyperpolarizing pyramidal cells directly. Similar to recent reports, we found that the former produced more effective and widespread silencing than the latter [27,30]. When inhibitory neurons were stimulated with ChR2 during testing, memory retrieval was significantly impaired. In contrast, when the same neurons were activated with the excitatory DREADD hM3Dq, retrieval was not affected. This dissociation was not due to differences in inhibition, as both manipulations activated interneurons throughout the dorsal hippocampus and reduced excitation (as indexed by c-Fos expression). Therefore, we hypothesize that the retrieval deficit caused by ChR2 stimulation is due to an immediate reduction in hippocampal activity that does not provide enough time for other brain regions to compensate. Stimulation of DREADDs, on the other hand, produces a gradual loss of excitation in the hippocampus that can take 10-30 minutes to reach asymptote [32,36]. This appears to be a sufficient amount of time for extra-hippocampal structures to become engaged and express context fear.

A similar difference between acute and prolonged hippocampal inactivation was observed during remote memory retrieval [18]. In that case, it was assumed that cortical regions could express context fear during prolonged silencing because systems consolidation had already taken place. However, we demonstrate that a similar dissociation exists 1-day after learning. It is possible, therefore, that cortical regions are able to express both recent and remote context fear provided they have enough to compensate for the loss of the hippocampus. It should be noted that the brain areas mediating memory retrieval in each of these situations may or may not be the same. The ACC is important for the expression of remote context fear while the PFC, entorhinal, perirhinal, postrhinal and retrosplenial cortices contribute to recent and remote memory retrieval [12,21,37–40]. The ventral hippocampus is also likely to play an important role as it projects to the amygdala and influences the expression of context fear [4,41–43].

The type of context memory that is expressed during prolonged inactivation may differ from the one that is retrieved in control animals. For example, context memories retrieved without the dorsal hippocampus are often less detailed and precise [16,44,45]. This possibility can be addressed in future work by using discrimination procedures to examine memory specificity [46]. However, it is important to note that damage to the dorsal hippocampus impairs retrieval on the same context tests that were used in the current experiments [6,7,22,23]. In the case of lesions, the deficits could result from consolidation impairments because memory is tested several days after surgery. The retrieval deficits could also be caused, in part, by damage to distal structures. Excitotoxic lesions of the hippocampus, for example, cause a substantial loss of cortical tissue that may prevent other brain regions from compensating during memory tests [47]. Consistent with this idea, patients with hippocampal damage exhibit episodic memory impairments that are strongly correlated with tissue loss in distal structures [48].

In contrast to prolonged inactivation, we found that rapid optogenetic silencing of the dorsal hippocampus produced significant deficits in memory retrieval. Given than other brain regions are able to express context fear, this result suggests that optogenetic inactivation somehow prevents them from doing so. One possibility is that acute silencing impairs retrieval because memory is tested at a time when activity is disrupted both in the hippocampus and in distal structures. For example, optogenetic perturbations have been shown to change local activity and simultaneously disrupt the function of distal brain regions [49,50]. These distal effects could prevent other areas from compensating for the loss of the hippocampus during memory retrieval. In contrast, when silencing occurs over an extended period, other brain regions have an opportunity to recover by the time memory is tested [49].

Consistent with a recent studies, we found that direct hyperpolarization of pyramidal cells produces less inhibition than stimulation of interneurons [27]. We hypothesize that restricted hippocampal silencing is the reason that some of our acute manipulations did not affect the retrieval of context fear. However, other groups showed that hyperpolarizing opins like NpHR can impair memory retrieval, although the extent of hippocampal inhibition was not reported [18]. It is likely the case that procedural differences between these studies affected the amount of silencing and/or the sensitivity of behavioral tests to hippocampal dysfunction.

Surprisingly, we found that some inhibitory manipulations (CaMKII-ArchT and hSyn-hM4Di) produced off-target effects that increased the total amount of hippocampal activity. When this occurred, memory expression was severely impaired. In addition, the deficits looked similar to those observed when pyramidal cells were activated with ChR2 or hM3Dq. In the case of CaMKII-ArchT, we hypothesize that the prolonged stimulation period (3-min.) increased excitatory activity by altering internal and/or external pH [34,35]. Shorter stimulation periods are recommended as they have been found to reduce these effects [33]. Reduced expression levels may also help as widespread increases in excitatory activity were not reported in previous ArchT studies that confined silencing to “engram cells” [3,51,52]. In order to properly interpret the effects of targeted optogenetic and chemogenenic manipulations on behavior it is clearly important to determine how they affect network activity [49,50]. In many cases, these effects go undetected because efficacy is confirmed *in vitro* and only in cells that express the protein of interest.

In summary, the current results demonstrate that the dorsal hippocampus is not required for the retrieval of recently acquired context fear memories. Other brain regions are able to retrieve this information provided they have adequate time to compensate and are not disrupted by artificial increases in hippocampal activity. These conclusions are illustrated in **Figure 7**.

**Figure 7.**
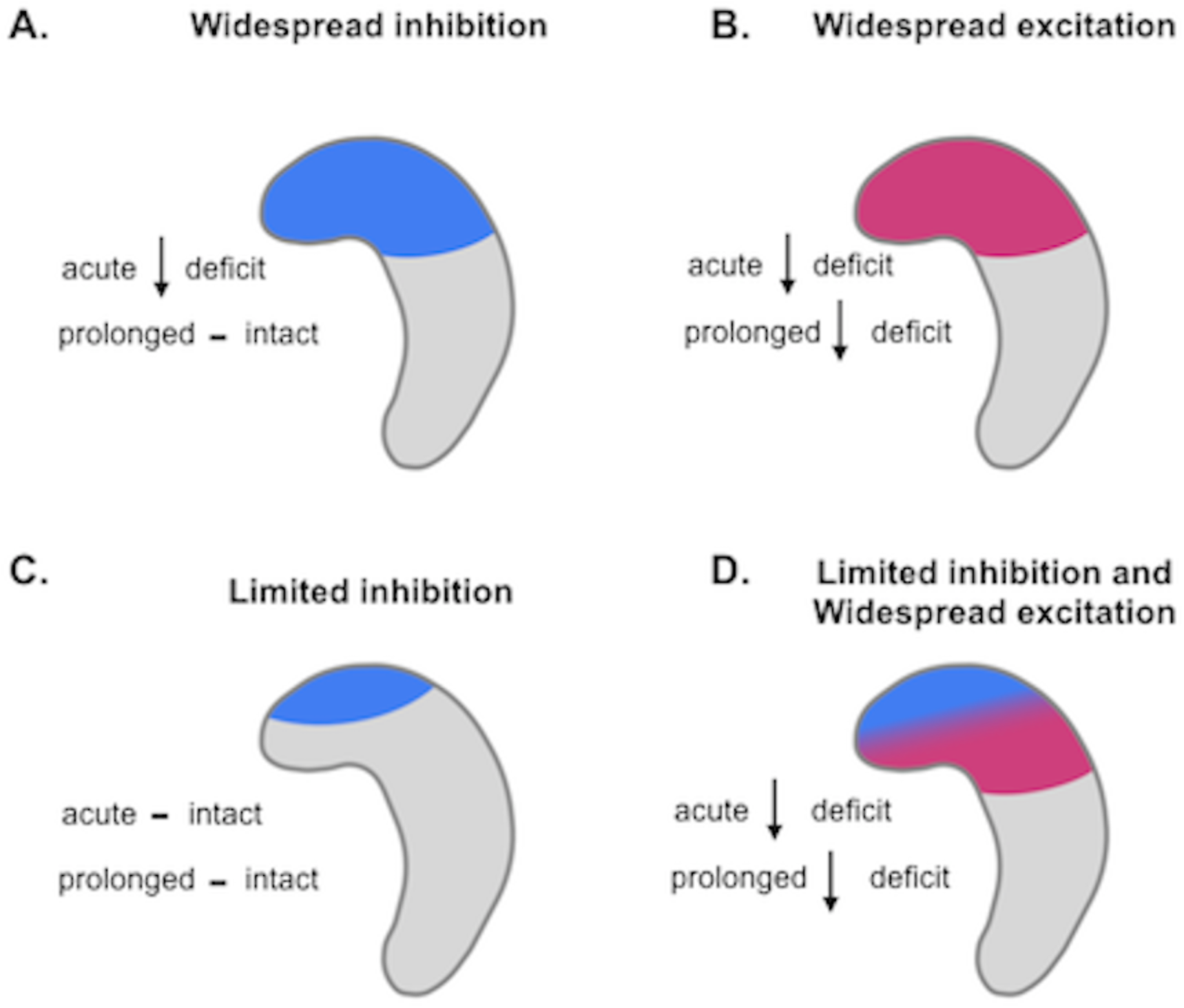
Manipulation of dHPC activity and the effects on memory retrieval. A. Widespread inhibition of excitatory activity throughout dHPC affects memory retrieval when inhibition is acute, but not prolonged. B. Widespread excitation (acute or prolonged) of dHPC impairs memory retrieval. C. Limited inhibition (acute or prolonged) of excitatory activity in dHPC does not affect memory retrieval. D. Limited inhibition accompanied by widespread excitation of dHPC (acute or prolonged) impairs memory retrieval.

## Materials and Methods

### Subjects

All procedures were approved by the Animal Care and Use Committee at UC Davis. Mice were maintained on a 12/12-light/dark schedule and were given free access to food and water. Animals were group-housed until surgery, after which they were single-housed for the remainder of each experiment. Surgeries were performed at 8 weeks of age, with behavioral experiments beginning at approximately 10-12 weeks of age. Both C57/B6J and F1 hybrids (C57/B6J:129SvEv) were used for these experiments, as described below.

### Stereotaxic Surgeries

All viruses were infused into the dorsal hippocampus (dHPC) (−2.0mm AP; +/− 1.5mm ML; −1.25mm DV) using a MicroSyringe pump controller at 2nL/second. For DREADD experiments, animals received 1µl bilateral infusions into the dHPC of *AAV5 CaMKII-hM4Di-IRES-mCitrine* (2.7×10^12 virus molecules/mL) (UNC Vector Core), *AAV5 hSyn-hM4Di-IRES-mCitrine* (1.12×10^12 virus molecules/mL) (UNC Vector Core), *AAV5 CaMKII-hM3Dq-IRES-mCitrine* (3.75×10^14 or 3.75×10^13 virus molecules/mL) (Addgene plasmid #50466, a gift from Bryan Roth, assembled and packaged with AAV5 helper plasmid by the UC Davis Molecular Construct and Packaging core facility. The virus was gradient purified, checked by SDS-PAGE and titered using qRT-PCR by standard methods [53], or *AAV5-hDlx-GqDREADD-dTomato* (3.15×10^15 virus molecules/mL) (Addgene plasmid # 83897, a gift from Gordon Fishell, assembled and packaged with AAV5 helper plasmid by the UC Davis Molecular Construct and Packaging core facility. The virus was gradient purified, checked by SDS-PAGE and titered using qRT-PCR by standard methods [53]. Dlx-enhanced viruses expressed red fluorophores (dTomato/mCherry) instead of green (mCitrine/eGFP/eYFP) due to commercial availability. We have successfully used AAV5 serotype viruses in the past and continued to use them for all DREADD experiments. For channelrhodopsin optogenetic experiments, animals received 0.25µl bilateral infusions of *AAV9 CaMKII-ChR2-eGFP* (8.96×10^11 virus molecules/mL) (Penn) or *AAV9 CaMKII-eGFP* (3.49×10^11virus molecules/mL)(Penn). For all other optogenetic experiments, animals received 0.5 µl bilateral infusions of *AAV5 CaMKII-ArchT-eGFP* (5.2×10^11 virus molecules/mL)(UNC), *AAV5 CaMKII-eYFP* (5.3×10^12 virus molecules/mL)(UNC), *AAV9 CaMKII-NpHR3.0-eYFP* (2.5×10^13 virus molecules/mL)(Penn), *AAV9 CaMKII-eGFP* (3.45×10^12 virus molecules/mL)(Penn) or *AAV5-hDlx-ChR2-mCherry* (7.6 ×10^14 virus molecules/mL) (Addgene plasmid # 83898, a gift from Gordon Fishell, assembled and packaged with AAV5 helper plasmid by the UC Davis Molecular Construct and Packaging core facility. The virus was gradient purified, checked by SDS-PAGE and titered using qRT-PCR by standard methods [53].

Following virus infusion, the injection needle remained in place for five minutes. The scalp of DREADD experimental animals was closed with surgical glue (Vetbond). For optogenetic experiments, fibers were implanted into the dHPC. Optic fibers were constructed and polished as previously described [54]. Briefly, a 200um diameter optic fiber (Thorlabs) was stripped and inserted into a plastic ferrule (PlasticOne). The convex side of ferrule was polished and the optical fiber scored with a ruby knife to extend 1.2 mm from the tip. Fibers were implanted into the dorsal hippocampus (−2.0mmAP; +/− 1.5mm ML; −1.1mm DV). Following implantation, the skull was scored with a surgical blade and covered in C&B Metabond (Parkell). A dental acrylic headcap (Harry J. Bosworth Company) was constructed to hold the fibers in place and seal the incision. Animals recovered for two weeks prior to the beginning of behavioral experiments to allow for recovery and sufficient receptor/opsin expression.

### Behavioral Apparatus

The contextual fear conditioning equipment used in all experiments was previously described [52,55]. Briefly, mice were trained and tested in conditioning chambers (30.5 cm × 24.1 cm × 21.0 cm) located within sound-attenuated boxes. The chambers consisted of a stainless-steel grid floor with overhead LED lighting (providing broad spectrum light). Sessions were recorded with a scanning charge-coupled device video camera (Med Associates). The chamber and drop pan were cleaned with 70% ethanol before each behavioral session. Detailed training procedures are provided below for specific optogenetic and DREADD experiments. Memory was assessed the following day by placing the mice in the context and measuring the freezing response. Freezing measurements were made using the automated Video Freeze System (Med Associates) as previously described [56].

### DREADD Behavioral Procedures

C57/B6J:129SvEv mice were ordered or bred in-house (Taconic). 8 groups were used in these experiments: CaMKII-hM4Di-CNO (3m, 3f); CaMKII-hM4Di-Vehicle (3m, 3f), hSyn-hM4Di-CNO (3m, 3f), Syn1-hm4Di-Vehicle (3m, 3f), CaMKII-hM3Dq-CNO (5m), CaMKII-hM3Dq-Vehicle (5m), Dlx-GqDREADD-CNO (7m, 3f), Dlx-GqDREADD-Vehicle (6m, 3f). After recovery, animals were handled for 5 minutes a day for five days prior to contextual fear conditioning for habituation to the experimenter. During training, following a three-minute baseline period, animals were given three two-second 0.4mA shocks with 60 second inter-trial intervals. The following day, animals were given IP injections of either 0.5 (CaMKII-hM3Dq-hM3Dq) or 5mg/kg (all other groups) CNO (Toronto Research Chemicals) (10mg dissolved in 100μL DMSO, diluted to 0.1 or 1mg/mL CNO in 0.9% saline) or vehicle (1% DMSO in 0.9% saline), one hour prior to re-exposure to the fear conditioning chamber. Animals were tested in the absence of shock for 30 minutes freezing behavior was analyzed during the first 12 minutes of the session.

### Optogenetic Behavioral Procedures

C57/B6J: mice used for these experiments were ordered or bred in-house (Taconic). Nine groups were used for these experiments: CaMKII-NpHr3.0 (3m, 3f), CaMKII-eGFP (3m, 3f), CaMKII-ArchT (3m, 3f), CaMKII-eYFP (3m, 3f), CaMKII-ChR2 20Hz (3m, 2f), CaMKII-ChR2 4Hz (4f, 1m), (CaMKII-eGFP (3m, 2f), Dlx-ChR2 (8f, 9m). Animals were handled for 2-5 minutes a day for five days prior to contextual fear conditioning. During handling, animals used for these studies were also connected to the optic fiber cable. Animals were placed in a Med Associates fear conditioning chamber with fiber implants connected to a splitting 200um optic fiber (Doric lenses). The fiber was attached to the conditioning chamber through a rotating commutator (Doric lenses) and coupled to a 473/532/561 nm 200 mW solid-state laser diode (OEM laser systems) with 15 mW output. Prior to each experiment, laser output intensity was measured with an optical power meter (Thorlabs). Stereotaxic coordinates were used to place optic fibers directly above dorsal CA1, directing laser stimulation to this region. Following a three-minute baseline period, animals were given four 2s 0.75mA shocks with 60-second inter-trial intervals. The following day, animals were returned to the chamber for a 12-minute testing session. Using Doric studio to trigger the laser, mice underwent laser stimulation twice for 3 minutes over the testing session. The session began with a 3-minute baseline period, then the laser (473nm (ChR2), 532nm (ArchT), 561nm(NpHR3.0), was turned on for 3 min (20Hz, 15ms pulses: ChR2; continuous stimulation NpHR3.0/ArchT) at 10 or 15mW. This was repeated once. Dlx-ChR2 animals underwent an additional testing session 24 hours later consisting of 12 minutes of laser stimulation.

### Tissue Collection and Immunohistochemistry

Ninety minutes following re-exposure to the chamber (DREADD experiments) or first laser on epoch (optogenetic experiments), animals were perfused using ice-cold PBS and 4% paraformeldahyde (PFA). Brain tissue was collected and stored overnight in 4% PFA. Slices were taken at 40nm using a Leica Vibratome. Tissue was stored in slice storage solution (100mL 10x tris-buffered saline, 300mL ethylene glycol, 300mL glycerol, 300mL dH_2_O) at −20°C until staining. Slices were washed 3 × 5 minutes in PBS, stained overnight in rabbit anti-cFos (1:5000, Millipore ABE457) in donkey blocking buffer. Slices were washed in 3 × 5 minutes in PBS, counterstained in biotynilated donkey anti-rabbit (Jackson) (1:500) for 1 hour, washed 3 × 5 minutes in PBS, stained with Cy3/Cy5 (Jackson) (1:500) for 45 minutes, washed in 3 × 5 minutes PBS and counterstained for 10 minutes with a DAPI nuclear stain (1:10000). Slices were mounted on SuperFrost slides with VectaShield mounting media.

### Microscopy and Cell Counting

Three to four coronal sections surrounding the −2.0 AP coordinate of each mouse were selected for c-Fos quantification. For optogenetic experiments, tissue was selected from beneath the fiber tip. Tissue slices were scanned in a 35um z-stack using an Olympus Slide Scanner at 20x magnification. Exposure times per channel (DAPI, FITC, TRITC) for each animal were set under saturation. Images were further cropped to the intermediate dCA1 for analysis. Fluorescent images were imported into ImageJ in grayscale and separated by channel. Cell counts were obtained using the multipoint tool on ImageJ with the experimenter blind to groups. An estimate of total dorsal CA1 cells per section was calculated by using 3D Object Counter in ImageJ and dividing the obtained volume by an average single nucleus volume for the group. Approximately 500 cells were counted from each slice, and 6 hemispheres were counted from each animal (∼ 3,000 cells). For Dlx experiments, c-Fos data quantified represent total c-Fos expression in pyramidal (c-Fos total neurons – cFos+/Dlx+ neurons) or inhibitory (c-Fos+/Dlx+) neurons per volume. Neuronal activity in inhibitory neurons was also quantified in inhibitory neurons (Dlx+ in the stratum oriens, stratum radiatum and stratum lacunosum-moleculare) at anterior (−1.2 to −1.5 AP), intermediate (−2.5 to −2.8 AP) and posterior (−3.2. to −3.5 AP) hippocampal regions. Anterior to posterior Dlx+ cell quantification was performed in single-plane images, and cell counts shown as number of c-Fos+ cells per ∼.45mm^2^.

### Quantification and Statistical Analysis

Freezing data during memory testing were analyzed using repeated measures ANOVA with p values set to 0.05. Post-hoc comparisons were made using Fisher’s LSD. All interactions and main effects are reported within results. Cell counts were normalized to control groups and analyzed using a t-test with p value set to 0.05. All data are represented as mean +/- SEM.

### Excluded data

For optogenetic experiments, subjects with poor fiber placement (defined as fiber not penetrating cortex or anterior/posterior from viral expression) were excluded from both behavioral and c-Fos analysis. For c-Fos analysis, some brain tissue was damaged and excluded from cell counts (1 CaMKII-hM4Di (CNO); 2 CaMKII-eGFP (NpHR control group); 1 CaMKII-eGFP (ChR2 control group); 1 CaMKII-eYFP (ArchT control group).

### Figures

Training/testing paradigm images (**Figures 2-3**) were created using *BioRender*. All data were plotted and analyzed using *GraphPad Prism 8*.

## Notes

#### Summary of Updates

Made some additional edits to the manuscript and uploaded the supplemental pdf.

