## Supplemental Figure 1 for "Recently formed context fear memories can be retrieved without the hippocampus"

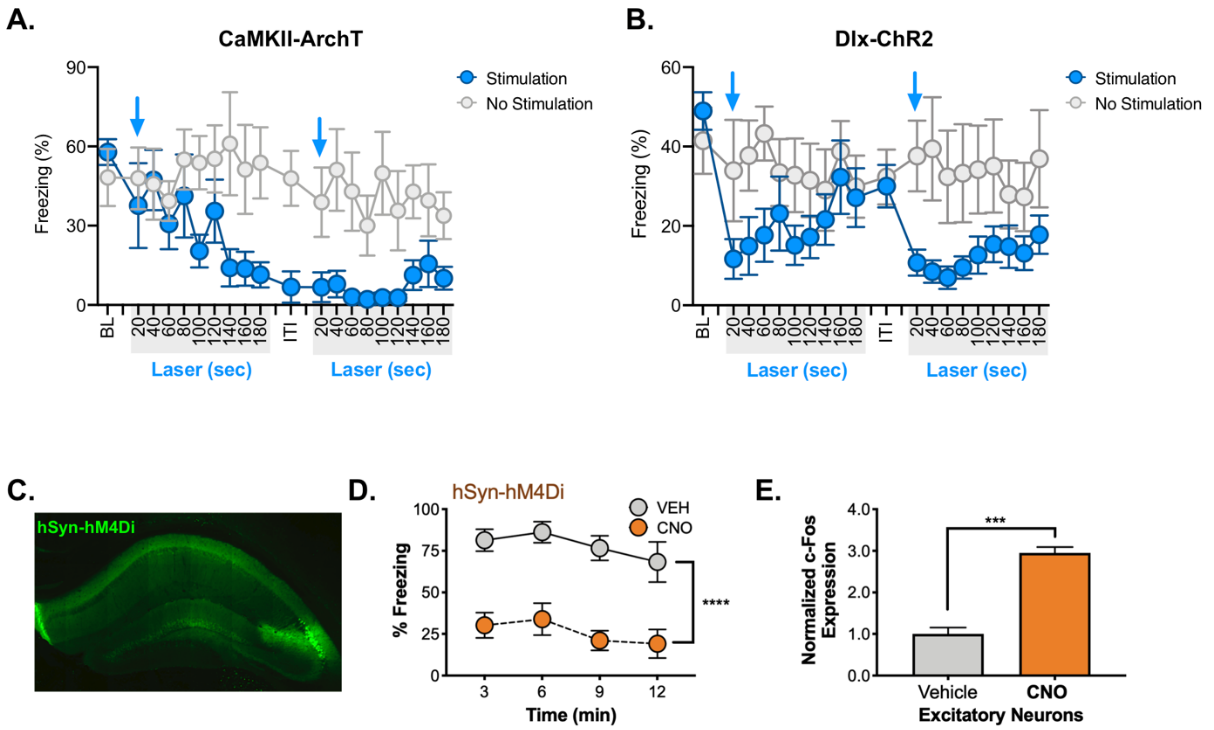


**Supplemental Figure 1**. A. 20 second bins of freezing activity in ArchT (blue) vs. control animals (gray) during laser ON periods. ~140s into laser stimulation, ArchT animals ceased freezing, this was not recovered over the remainder of the testing period. Blue arrows denote onset of laser stimulation. B. 20 second bins of freezing activity in Dlx-ChR2 laser-stimulated (blue) vs. control animals (gray) during laser ON periods. Dlx-ChR2 animals ceased freezing during laser onset (blue arrows), but freezing recovered during laser OFF periods. C. hSyn-hM4Di virus expression (green) in dHPC. D. During the 12-minute testing period, hSyn-hM4Di CNO-treated animals froze less than vehicle treated animals. E. c-Fos expression was increased in excitatory neurons of dCA1 in CNO-treated animals. Data are expressed as mean +/- SEM.
